# Costs of antibiotic resistance genes depend on host strain and environment and can influence community composition

**DOI:** 10.1101/2023.09.07.556771

**Authors:** Huei-Yi Lai, Tim F. Cooper

## Abstract

Antibiotic resistance genes (ARGs) is a major contributor to increasing levels of antibiotic resistance in clinical and agricultural settings. ARGs are strongly selected in environments containing corresponding antibiotics, but it is less clear how ARGs are maintained in environments where antibiotic selection might be weak or sporadic. In particular, few studies have directly estimated the effect of ARGs on host fitness in the absence of direct selection. To the extent that ARGs impose costs, it is not clear if these are fixed or might depend on the host strain, perhaps marking some ARG-host combinations as low-cost reservoirs that can act to maintain ARGs in the absence of antibiotic selection. We quantified the fitness effects of six ARGs in each of 11 diverse *Escherichia* spp. strains in two environments. While three ARGs *(bla*_TEM-116_, *cat*, and *dfrA5*, encoding resistance to β-lactam antibiotics, chloramphenicol, and trimethoprim, respectively) imposed an overall cost, all ARGs had an effect in at least one host strain, reflecting a significant ARG effect-by-strain interaction effect. A simulation model predicts that these interactions cause the ecological success of ARGs to depend on available host strains, and, to a lesser extent, for successful host strains to depend on the ARGs present in a community. Together, these results indicate the importance of considering ARG effects over different host strains, especially the potential of reservoir strains that allow resistance to persist in the absence of direct selection, in efforts to understand resistance dynamics.

**Importance:** Antibiotic resistance is a major and increasing public health concern. Resistance occurs through a variety of mechanisms but commonly involves bacterial strains acquiring antibiotic resistance genes (ARGs) encoded by mobile elements. It is obvious and well-documented that ARGs will be selected in bacteria that are exposed to the antibiotics they confer resistance to. ARGs can also confer costs to bacteria—in environments that do not contain antibiotics, these costs can lead to the loss of ARGs. We show that ARG costs can be significant and that they depend on the host bacterial strain and the environment in which strains are grown. This dependence creates host-environment refuges for many ARGs, allowing them to be maintained in the absence of direct selection.

## Introduction

Widespread use of antibiotics has led to selection for antibiotic-resistant bacteria, which has been identified as a major global health concern [1]. Resistance can derive from spontaneous mutations that generally reduce affinity or access of antibiotics to cellular targets and from antibiotic resistance genes (ARGs) that typically act to detoxify or export antibiotics from cells [2]. The link between antibiotic exposure and selection of resistance is clear: ARGs tend to rise in frequency shortly after widespread use of the cognate antibiotics and ARGs of recent origin are associated with host bacteria that can cause infection in humans or domestic animals [3–6]. Even exposure to low levels of antibiotics can confer a significant advantage to resistant cells, potentially allowing selection in a wide range of environments [7].

The benefit conferred by ARGs to host cells during antibiotic exposure is, however, only one factor determining their overall success [8–11]. Of 22 ARGs for which a recent origin could be inferred, most were from species with human or domestic animal association [5]. This association is linked to antibiotic exposure, supporting the idea that direct selection for resistance is important for the success of ARGs [12]. However, many species that are associated with humans or domestic animals, will nevertheless transit through environments with low antibiotic concentrations. For other ARGs, no originating host species can be inferred, consistent with them originating in poorly characterized environmental species [5]. These considerations underlie the view that any costs associated with encoding ARGs will play an important role in determining their overall success in bacterial populations [10,13,14].

Costs of antibiotic resistance have been examined in some detail for spontaneously occurring resistance alleles [15]. In one experiment, costs associated with rifampicin resistance were sufficient to drive selection for frequent reversion to sensitivity in a mouse infection model [16]. In other cases, costs drive the selection of secondary mutations that compensate for the costs of resistance allowing the resistance phenotype to persist with relaxed antibiotic-driven selection (reviewed in [13,15]. Spontaneous resistance often occurs through mutations in essential genes encoding products targeted by the relevant antibiotic [17]. Changes to these products will generally alter their structure in some way that reduces the strength of antibiotic interaction and are likely to have some deleterious effect on the original function [14]. By contrast, less work has been done examining effects to host cells of expressing introduced ARGs.

Studies have shown fitness costs to hosts that carry ARG-encoding plasmids but estimates of the effect of ARGs themselves are often indirect and differences in experimental designs make comparisons difficult. One meta-analysis found that the costs associated with plasmids generally increase with the number of encoded ARGs [14]. That study found some support for interactions between resistance genes affecting their cost, but it was not possible to identify costs of specific ARGs or to determine if those costs depend on the host strain in which they are assessed. Studies estimating the effect of specific ARGs include analysis of the *tetAR* operon, encoding tetracycline resistance, that was estimated to impose a cost of around 1% in rich medium [18,19]. Similarly, β- lactamases, a large class of resistance genes that confer resistance to one or more of the β-lactam antibiotics, have been shown to decrease host fitness, though different variants have different effects in different hosts [20–23]. In at least some cases, however, ARGs, including *tetAR*, not only do not confer a cost but can provide a benefit to host cells, underlining the importance of understanding potential variation in costs across different hosts [24–26].

The possibility that ARGs have different costs in different hosts is especially important to understand because of the potential for costs of ARGs to be transmitted between different hosts through an association with horizontally mobile elements (reviewed in 12). Studies of mutation fitness effects, including some conferring antibiotic resistance, often find significant differences depending on the host background [27–32]. For example, although not isolating the effect of the ARG itself, a plasmid encoding the *bla*_OXA-48_ resistance gene conferred an overall cost across a range of host strains, but with a range between ∼20% cost and ∼20% benefit [33]. Simulations predict that this high variation can increase plasmid, and, therefore, ARG, persistence in a multi-member community because hosts in which it has low or no cost can serve as reservoirs during periods when the plasmid is not directly selected [33]. These results make it clear that any attempt to understand the dynamics of ARGs needs to account for the possibility of differential effects in different genetic, and likely, environmental, contexts.

We separately cloned six ARGs representing four resistance classes into a control plasmid and estimated their fitness effect in each of 11 genetically diverse *Escherichia* spp.. We found that costs were diverse across ARGs and strains, and that they interact so that different strains have the potential to act as no-cost reservoirs for different ARGs. Simulations predict that the magnitude of ARG effects are sufficient to cause both the presence of different ARGs to influence which strains can persist in a community and the availability of strains to determine which ARGs can be maintained. Our study makes clear that ARGs should not be simply considered as costly (or neutral). Instead, effect depend on the host strain in which they are assessed. This dependence is important to consider in attempts to predict the ability of ARGs to persist in antibiotic-free environments.

## Materials and Methods

### Selection of candidate ARGs

The ARGs used in this study were chosen based on their prevalence in accessible *E. coli* genomes and their availability in our laboratory [34]. The 2022 release of the CARD Prevalence, Genomes and Variants dataset gives the prevalence of our chosen ARGs among sequenced *E. coli* genomes as: *aadA* (ARG type = 15.3%; specific ARG allele = 1.73%), *cat* (5.3%, 4.1%), *dfrA5* (19.4%, 1.8%), *bla_TEM-116_* (22.7%, 0.07%), *bla_CTX-M-15_* (17.1%, 5.83%) and *bla_SHV12_* (0.59%, 0.04%), identifying them as clinically important ARGs.

### Construction of ARG-plasmids and bacterial strains

Focal ARGs were originally cloned into pUA66, which contains a promoterless *gfp* [35]. However, we found leaky expression of GFP in this vector (data not shown). To avoid any influence of GFP expression on fitness measurements, we removed the GFP by amplifying the pUA66 backbone (using pUA_minusFP_F: 5’atgtccagacctgcaggcatg; pUA_minusFP_R: 5’ggatccatcgaggtgaagacg) and circularizing the product with a bridging oligonucleotide (5’cgtcttcacctcgatggatccatgtccagacctgcaggcatg) using the NEBuilder® HiFi DNA Assembly (NEB) kit. The resulting plasmid, pmFP, retains the kanamycin resistance gene originally present on pUA66. The six ARGs were amplified from template sources as listed in Table S1 and separately added into the pmFP vector (amplified using primers: pUA66_F: aataggcgtatcacgagg and pUA66_EcoRI_R: gaattcatggtttcttagacgtcgg) using NEBuilder® HiFi DNA Assembly. All PCR reactions used Q5® High-Fidelity polymerase (NEB) unless otherwise specified. When sequencing cloned ARGs we found that the *bla_TEM-116_* gene in pTarget, which was used as a template, has a non-synonymous mutation (Q274R) relative to the reference *bla_TEM-116_* sequence (NCBI accession: U36911). To distinguish between these genes we identify the variant used here as *bla_TEM-116_\**.

The six constructed pmFP::ARG plasmids, and the reference pmFP backbone vector, were each used to transform one *E. albertii* and 10 genetically divergent *E. coli* strains (Fig. S1) following the method outlined in ref. [36]. After transformation, cells were recovered at 37 ℃ for one hour before being plated on lysogeny broth (LB) supplemented with 50 μg/ml kanamycin to select plasmid containing cells. One strain, H305, was resistant to ampicillin, chloramphenicol, and trimethoprim, antibiotics. Omitting H305 makes no qualitative difference to the interpretation of any analysis we present, though we note that H305 was one of two strains in which no ARG conferred a significant effect on fitness and one of three strains in which no ARG was costly. We leave it in the presented analyses because already resistant strains can clearly serve as hosts to ARG-encoding plasmids in natural communities and because it is reasonable to think that additional copies of a given resistance gene can impose some additional fitness effect.

### Fitness competitions

To measure the fitness effect of each ARG, we performed a series of competition assays. The fitness effect of an ARG was estimated as the fitness difference between cells carrying an ARG plasmid (W_ARG_) and cells carrying the control pmFP plasmid vector (W_Vec_). The fitness of ARG and vector plasmid-carrying cells was estimated indirectly by competing each against reference cells carrying a GFP expressing plasmid (pUA66-_PrpsL_GFP).

Cells carrying an ARG plasmid, pmFP, or pUA66-_PrpsL_GFP, were inoculated in 200 μL of Davis-Mignioli broth supplemented with 250 μg/ml glucose (DM250) and 50 μg/ml kanamycin from frozen stocks and cultured at 37°C overnight. The overnight cultures were diluted 1:100 in 200 μL of fresh DM250 and incubated for a 24-hour growth cycle. Cultures were grown over two additional daily growth cycles with 1:100 dilution between each cycle to condition them to the antibiotic-free medium, then used to set up competition assays. Five microliters of culture of each competitor – reference cells with pUA66-_PrpsL_GFP and either cells with an ARG plasmid or with the control plasmid – were mixed in 40 μL of fresh DM250. Forty microliters of the mix was added to phosphate buffered saline (PBS) and placed on ice or fixed with formaldehyde before being assayed by flow cytometry to determine the proportion of reference (GFP+) and ARG or control plasmid-containing cells (Day 0). The remaining 10 μL of the mix was added to 190 μL of fresh DM250 and incubated for 24 hours to allow strains to compete. Following this competition, cultures were diluted 20-fold in PBS and assayed by flow cytometry (Day 1). This protocol was repeated to obtain ARG effect estimates in a competition environment supplemented with kanamycin, to which all plasmids confer resistance to. At least five independent fitness measurements were collected for each host strain-ARG combination. Construction of some combinations occurred later than others due to factors such as delays in constructing ARG vectors. For this reason some combinations have higher replication than others.

The fitness effect of an ARG was estimated as W_ARG_ / W_Vec_, where W_ARG_ = ln(ARG_Day1_ × 100 / ARG_Day0_) / ln(GFP_Day1_ × 100 / GFP_Day0_) and W_Vec_ = ln(Vec_Day1_ / Vec_Day0_) / ln(GFP_Day1_ / GFP_Day0_). ARG_Day1_, ARG_Day0_, Vec_Day1_ and Vec_Day0_ are the proportion of flow cytometry events without a GFP signal, and GFP_Day1_ and GFP_Day0_ are the proportion of events with a GFP signal. Day 1 proportions are multiplied by 100 to account for growth occurring during the competition. We note that a number of flow cytometry runs resulted in total event counts or counts of one competitor that were substantially different to mean estimates over all replicates of any given competition type. The most likely reason for this is that the natural isolate strains used here are subject to clumping that introduces variability in passage of cells through the cytometer flow cell. To reduce the effect of this variation we excluded competitions where the starting frequency of either competitor was estimated as more than 15% different than the target of 50%. This step removed a total of 810 of 4372 competitions. We emphasize that this filtering was performed without regard to the fitness estimated considering the change in frequency over the course of the competition. We also omitted a further 16 fitness estimates of one host strain collected in one experimental block. These estimates were more than 50% higher than a set of 205 estimates collected for the same strain in other experimental blocks, indicating that they were compromised by some experimental error.

### Assessing plasmid loss and compensation

In the antibiotic-free assay environment it is possible that plasmid loss could occur, which could influence fitness estimates through production of relatively higher fitness plasmid free sub-populations. To estimate the influence of plasmid-losing cells during the competition assay, monocultures of cells with the pUA66-_PrpsL_GFP plasmid were grown following the competition protocol and plated on non-selective media to allow identification of the proportion of plasmid containing cells. After three transfer cycles the frequency of cells that lost the plasmid was 0.004 (± 0.004 95%CI, n = 4). If we conservatively, consider this frequency to represent the fraction of cells incorrectly identified as being the ARG-plasmid containing competitor instead of cells that lost the reference pUA66-_PrpsL_GFP plasmid during a single day of competition, this would represent a fitness inflation of the ARG of only <0.1% over the true value. Moreover, plasmid-loss cannot easily explain differences in costs of different ARGs in the same host. Finally, we repeated fitness estimates in an environment supplemented with kanamycin, which ensures that only plasmid-containing cells are present. The overall magnitude of fitness effects in this environment were similar to those in a kanamycin-free environment, suggesting that plasmid loss is not influencing fitness estimates in the conditions used in our experiment.

Compensatory mutations that reduce the cost of an ARG could occur during our competition assays. In practice, however, they are very unlikely to significantly effect fitness estimates. For example, a compensatory mutation conferring a benefit of 15% (i.e., completely compensating the largest cost found in our experiment) starting at ten copies in populations of the size used in our experiment will reach a frequency of ∼0.0001 in 60 generations, a span that coves the growth of a freezer stock, competition acclimation, and the competition itself.

### Determination of copy number of the bla_TEM-1_ plasmid

The copy number of plasmids encoding the *bla_TEM-116_\** ARG was measured by quantitative PCR (qPCR) using the comparative Ct (ΔΔCt) method [37]. Plasmid carrying cells were cultured overnight in DM250 supplemented with 50 μg/ml kanamycin and DNA isolated using a Wizard Genomic DNA Purification Kit (Promega). Amplification of target genes was carried out using a SYBR Green based qPCR mix consisting of Q5^®^ High-Fidelity 2× Master Mix (NEB), SYBR Green, and relevant primers (at a final concentration of 0.25 μM). Between one and two ng of template DNA was used per 10 μl reaction. Reactions were performed using a Thermo Scientific PikoReal Real-Time PCR System with default settings. Primers were designed using PrimerQuest (Integrated DNA Technologies, Inc.) to amplify the chromosomal *dxs* gene, in order to determine the absolute Ct value of the bacterial chromosome (Ct_ch_; xs_qpcr_F1:cgagaaactggcgatcctta; dxs_qpcr_R1: cttcatcaagcggtttcaca), and the pUA66 plasmid encoded *aph(3’)-IIa* gene, to determine the Ct value of the plasmid (Ct_p_; pUA_Kan_qpcr_F1: ctcgtcaagaaggcgatagaag; pUA_Kan_qpcr_R1:cgttggctacccgtgatatt). The relative copy number of the *bla_TEM-116_\** plasmid to the chromosome (ΔCt) was calculated as 2 ^-(*ct_p_* - *ct_ch_*)^. The ΔCt of different bacterial strains was then normalized to the lab strain, REL606, to obtain the ΔΔCt value.

### Statistics, phylogeny construction and phylogenetic signal testing

The R statistical computing platform was used for all analysis and visualization [38]. The flowCore package [39] was used to analyze flow cytometry data and the lme4 and lmerTest [40,41] packages were used to run and evaluate mixed models. Environment, ARG and host strain were included as random effects in these models as noted in the text. We isolate the fitness effect of an ARG in each ARG-strain combination as the difference in fitness effect of a plasmid encoding the ARG to the fitness effect of the vector pmFP plasmid. To account for the comparison of each ARG to the same vector pmFP plasmid we use Dunnett’s tests implemented in the multcomp package, which control for many-against-one comparisons [42]. Core and accessory genomes of a collection of 96 environmental *E. coli* isolates were determined and used previously to build phylogenies [43]. These phylogenies were used to test for a phylogenetic signal of ARG effects using Pagel’s lambda and Blomberg’s K metrics as implemented in the phylosig function of the phytools package 1.0-3 [44].

### Simulation of host-ARG communities

Competitive fitness assays allow fitness of plasmid-containing strains to be measured and compared to a corresponding ancestral strain. It is not possible to extend this approach to compare the fitness of our distinct progenitor strains because competition between these strains may be affected by complex interactions, for example mediated by colicins or phage, that lead to non-transitivity in their relative fitness [45]. To estimate the fitness of all strains, we analyzed growth curves of progenitor strains to estimate their fitness relative to one another as determined only by differences in resource use [45]. The fitness of each strain-ARG combination was determined by analysing the growth curve of each progenitor strain to determine its baseline fitness and then altering that baseline to account for the fitness effect of each ARG measured in that strain (as measured by competition assays). Resulting fitness estimates of each strain-ARG combination were used to parameterize an individual based model that simulated dynamics in a community with strain-ARG fitness drawn form a normal distribution with mean and standard deviation taken from experimental estimates. We draw from a distribution of fitness values to account for the measurement error inherent in our estimates and note that it can mean that ARGs with a higher mean relative cost, but larger distribution width, can be more successful than more fit ARGs with a smaller distribution width. Simulations were initiated with equal numbers of each strain-ARG combination in a population of size 10^4^ individuals and propagated using a Wright-Fisher model for 200 generations. Strains and ARGs remaining above a threshold frequency of 0.01 in at least 20% of 10 replicate simulations were considered to have persisted in the community. Simulations were repeated separately omitting each strain and each ARG from the starting community to determine the effect of each individual strain and ARG in determining the final community composition. The model includes only fitness differences of strains integrated over a complete competition cycle. As such, feedbacks are not present and a single strain-ARG combination will eventually win in each simulated competition. This is not a realistic, scenario and we currently have experiments underway to examine interactions directly. Nevertheless, our model does clearly demonstrate that costs of ARGs are sufficient that they can be expected to effect competition outcomes.

## Results

### ARGs can impose costs and benefits in the absence of direct selection

We separately cloned six ARGs belonging to four distinct mechanistic classes into a low-copy number plasmid vector (Table 1). The fitness effects of each ARG was estimated in an antibiotic-free environment in each of 11 divergent *Escherichia* spp. host strains (Fig. 1). We found significant differences in the mean effect of the different ARGs (χ^2^ = 40.4, P < 0.001). Of the six tested ARGs, three conferred a significant overall change in fitness measured across all host strains: *bla_TEM116*_* conferred a 2.7% cost (±1.0% 95CI; Dunnett’s test P < 0.001), *cat* conferred a 1.8% cost (±1.2% 95CI; Dunnett’s test P = 0.01) and *dfrA5* conferred a 1.6% cost (±1.2% 95CI; Dunnett’s test P = 0.05). This result clearly indicates that ARGs can confer significant effects on fitness in the absence of direct selection for their resistance phenotypes. Moreover, the three ꞵ-lactamases had significantly different effects on host fitness, demonstrating that effects can differ even between ARGs with similar resistance profiles (χ^2^ = 32.1, P < 0.001).

**Figure 1.**
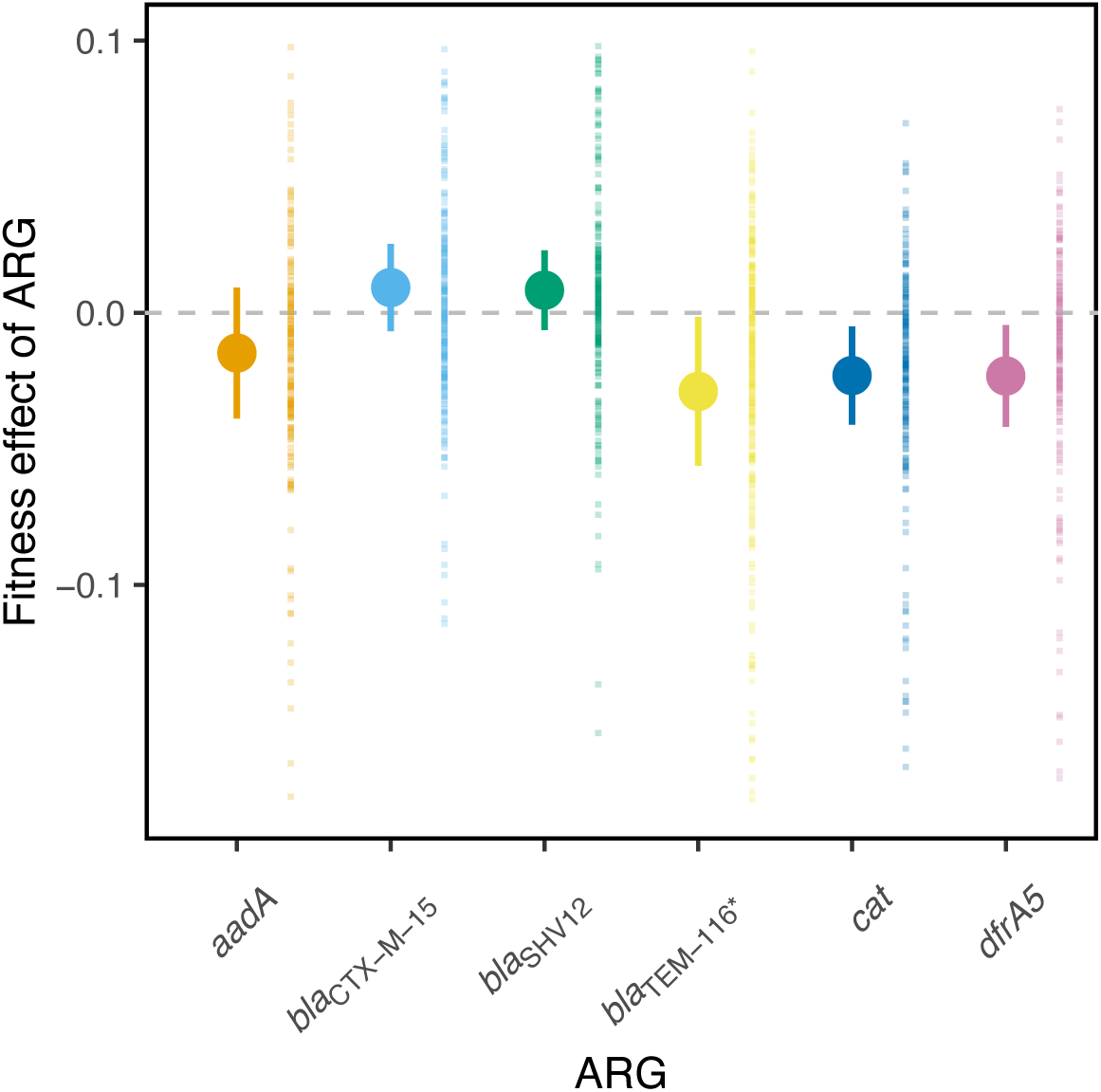
Fitness effect of each ARG across 11 *Escherichia* spp. host strains. For each ARG the solid symbol indicates mean fitness over all replicate estimates and error bars indicate 95% CI. Small background points indicate individual estimates used to calculate each mean. The horizontal dashed line indicates no fitness effect. n > 200 for each ARG (summed across all 11 host strains, see figure 2 for effects per host strain).

**Table 1.** ARGs used in this study.

| Gene | Enzyme type | Conferring resistance to: | Mechanism |
| --- | --- | --- | --- |
| *aadA* | Aminoglycoside adenylyltransferase | Aminoglycosides such as spectinomycin and streptomycin | Drug-modification |
| *bla<sub>TEM-116</sub>\** | Beta lactamase | β-lactam such as ampicillin | Drug-modification |
| *bla<sub>SHV12</sub>* | Beta lactamase | β-lactam such as ampicillin and cefotaxime | Drug-modification |
| *bla<sub>CTX-M-15</sub>* | Beta lactamase | β-lactam such as ampicillin and cefotaxime | Drug-modification |
| *cat* | Chloramphenicol acetyl transferase | Chloramphenicol | Drug-modification |
| *dfrA5* | Dihydrofolate reductase | Trimethoprim | Target replacement |

### Fitness effects of ARGs depend on the host strain

The results presented above consider the overall effect of each tested ARG over 11 host strains. To determine if effects differed between strains, we extended our model to include an ARG effect-by-strain interaction term. This extended model provided a significantly improved fit compared to the previous model, indicating a dependence of ARG effects on host genotype (χ^2^ = 94.9, P < 0.001). We found that all ARGs conferred a significant fitness effect in at least one strain (Dunnett’s test P < 0.05: *aadA* - 4 strains; *bla_CTX-M-_*_15_ - 2 strains; *bla_SHV12_* - 1 strain; *bla_TEM116*_*, *cat*, *dfr*A5 - 3 strains) (Fig. 2). In 11 of 15 cases significant effects were negative, with every ARG except *bla_SHV12_* imposing a cost in at least one strain. From the perspective of host strains, all except H305 and B354 had a significant change in fitness caused by at least one ARG (Dunnett’s test P < 0.05). The mean fitness difference between different ARG-strain combinations was substantially larger than the mean difference between ARG effects, indicating a potential for some strains to act as reservoirs for ARGs in antibiotic-free environments (ARG-Strain combinations mean difference = 2.5% (1.9 – 3.3% 95CI); ARG mean difference = 1.2% (0 – 2.8% 95CI)). Indeed, all of the six tested ARGs were neutral or conferred a benefit in at least one host strain.

**Figure 2.**
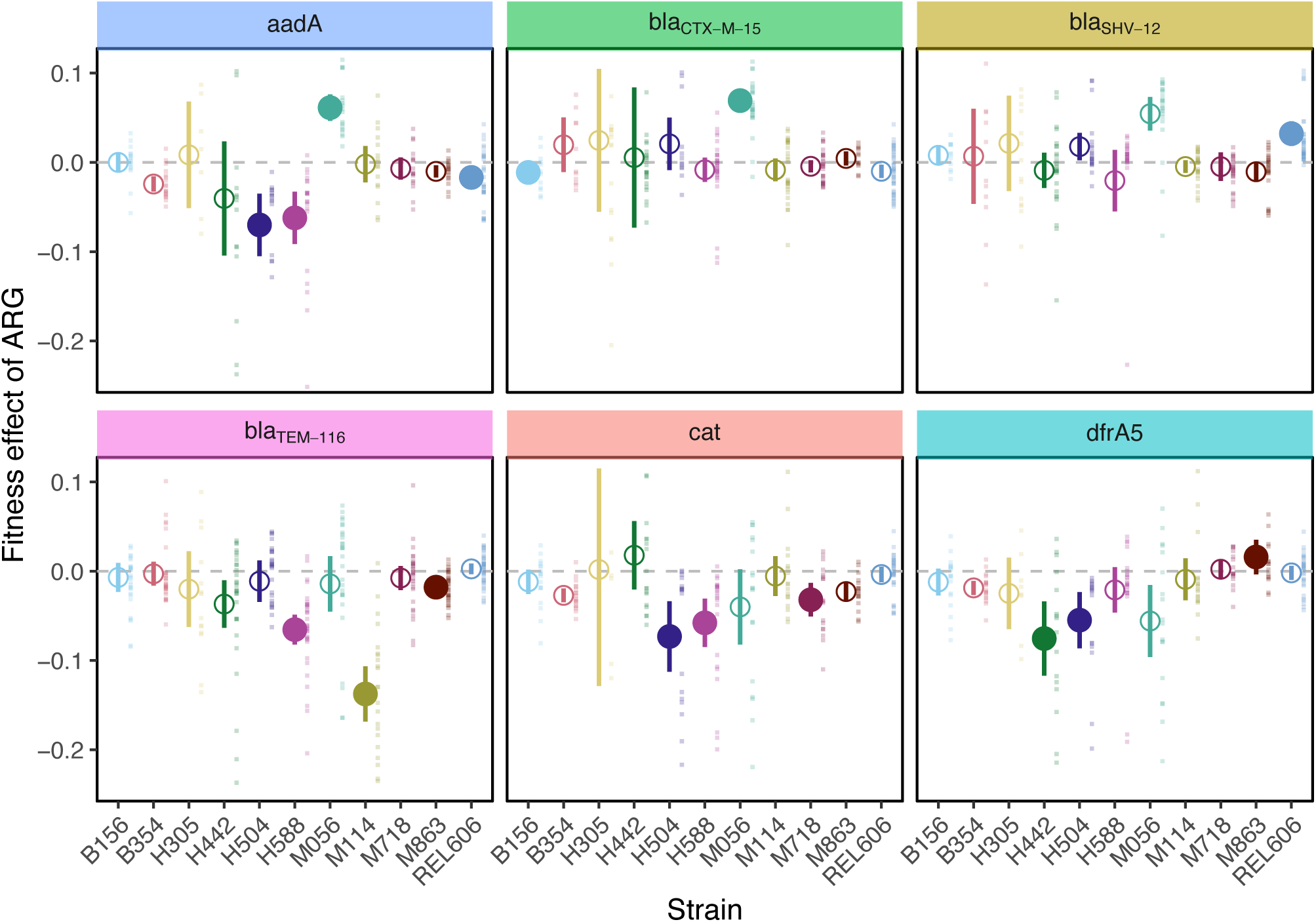
Fitness effect of ARGs in each host strain. For each host-ARG combination large symbols indicate mean fitness over all replicate estimates and error bars indicate 95% CI. Large symbols are filled if the mean ARG effect is different from 0 after correction from multiple comparisons using Dunnett’s test (M&M for details). Background points indicate individual estimates used to calculate each mean. The horizontal dashed line indicates no fitness effect. Background label panels are coloured to match the ARG color used in figure 1. n ≥ 5 for each host-ARG combination.

### Fitness effects of ARGs depend on the environment

Fitness estimates reported above were carried out in a minimal medium environment in the absence of any antibiotic. The focal ARGs were cloned into a vector that also included an additional ARG gene (*aph3’-II*, encoding resistance to kanamycin), giving us the opportunity to test if a secondary antibiotic might influence the physiology of strains in a way that alters the effect of non-selected ARGs. We note that kanamycin is an aminoglycoside antibiotic with some similarity to spectinomycin and streptomycin, substrates for the *aadA* ARG, but there is no detectable cross-activity [46]. When we measured the fitness effects of each ARG-strain combination in the same environment as used previously, but supplemented with kanamycin, we again found a significant interaction between ARG and host strain in determining fitness (χ^2^ = 47.0, P < 0.001)(Fig. S2). The fitness of all strains were significantly affected by at least one ARG and all ARGs except *aadA, cat* and *bla_SHV12_* significantly affected fitness of at least one strain. There was a significant effect of the environment when comparing fitness estimates made in the minimal medium and kanamycin supplemented environments (χ^2^ = 74.4, P < 0.001). Evidently, a general pattern of ARG fitness effects depending on strain background was common across the environments, although the exact nature of that dependence is different.

### Fitness effects of ARGs are not explained by plasmid copy number

A possible explanation for different effects of ARGs across host strains is that the plasmid they are encoded on has a different copy number in different strains. Different copy number can affect expression levels of plasmid-borne genes and, thus, any phenotypic effect of gene products. One well known example is the effect of *bla*_TEM1_ copy number on the degree of ampicillin resistance [47]. We used qPCR to determine the copy number of the *bla_TEM116*_* plasmid across our set of 11 host strains. This ARG showed the highest variation in fitness cost so represents a good test of the contribution of copy number differences to differences in fitness effects. We found that the relationship between *bla_TEM116*_* copy number and fitness cost was positive, but non-significant, in both the antibiotic-free environment and in the kanamycin-supplemented environment, indicating that differences in plasmid copy number are unlikely to represent a general mechanism of strain-specific ARG fitness effects (minimal: r = 0.28, P = 0.36; kanamycin-supplemented: r = 0.26, P = 0.41).

### ARG effects are not predicted by evolutionary relationships between host strains

Variation in ARG effects across host strains must derive from differences in their genotypes. To test if the dependence of ARG effect on host strain can be predicted through knowledge of the genetic relationship of strains we tested for a phylogenetic signal in the effect of each ARG. We estimated Pagel’s λ, a measure of the phylogenetic signal in a response variable, considering the fitness effect of each ARG across phylogenies derived from the core and accessory genomes of our host strains, excluding B156, which we do not have a genome sequence for [48]. In no case was any significant signal found. An alternative measure of phylogenetic signal, Blomberg’s K [49], gave a qualitatively consistent result (Table S2). With only 11 host strains, the power of individual tests is not high. However, the lack of signal given that our host strains are genetically divergent and that tests were repeated for each of our six ARGs, indicate, at least, that that fitness effects of the tested ARGs are not well predicted by the overall genetic similarity of diverse host strains.

### Dependence of ARG fitness effect on host strain predicts interdependence between host strain and ARG success

The above results demonstrate that a range of ARGs confer different fitness effects in different host strains. We predict that this interaction can cause the success of host strains and ARGs to be interdependent: the overall fitness of each ARG depends on available host strains and the overall fitness of each host strain depends on the ARGs it carries. A consequence of this dependence is a potential for some strains to act as low-cost reservoirs of ARGs, allowing these ARGs to persist when they would not otherwise be able to. It is also possible that the presence of some ARGs will affect the relative fitness of some strains allowing them to be maintained in a community when they would otherwise be outcompeted.

As a first step to examine implications of host-ARG fitness interdependence we simulated 200 generations of growth of a community started with equal numbers of all 66 host-ARGs combinations considered in our study. At the end of the simulation strains and ARGs remaining above a threshold frequency in at least 20% of replicate simulated communities were counted as having persisted (see Materials and Methods for details). We found that the final community contained one host strain, B354, and a single ARG, *bla_SHV12_* (Fig. 3A and B, top right squares). This reflects that these strain-ARG combinations had the highest fitness of any that we measured (Fig. 2 combined with baseline strain fitness estimates, see Materials and Methods for details). To determine the effect of strain dependent ARG costs on this result, we repeated simulations separately omitting each strain and ARG. In some cases, quite different final communities were produced (Fig. 3). For example, omission of the B354 strain from the starting community led to the H504 strain dominating. In this strain as many as three of the six focal ARGs remained present at the end of the simulation period, reflecting the relatively small and consistent effects of ARGs in the H504 host strain.

**Figure 3.**
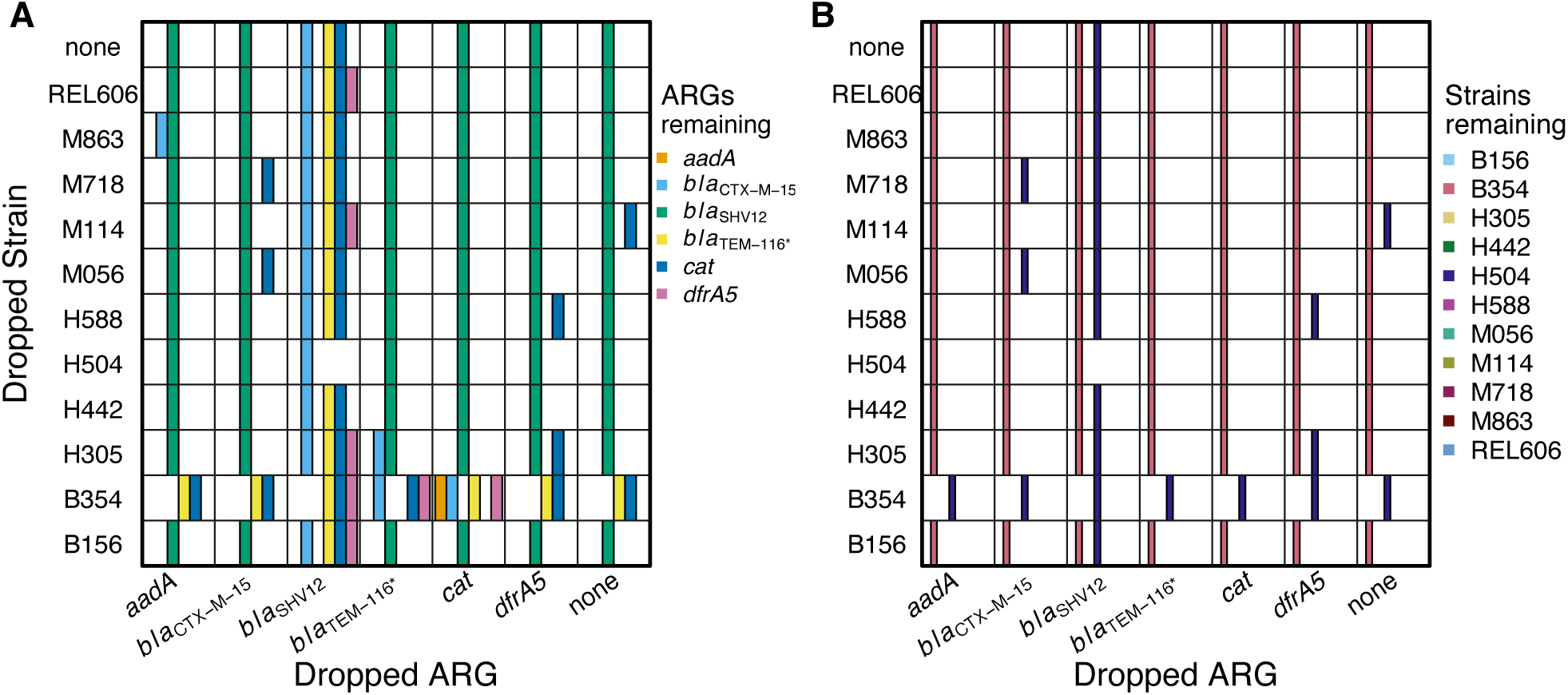
Host strain and ARG persistence during community growth in the absence of direct ARG selection. Communities consisting of all 66 host strain-ARG combinations (top right squares of each panel) and with omission of one strain (Dropped Strain axis) and/or one ARG (Dropped ARG axis) were simulated over 200 generations using fitness values that combined a baseline fitness of each strain with changes in fitness caused by addition of ARGs to that strain (details in Materials and Methods). Coloured bars indicate the ARGs (panel A) and host strains (panel B) remaining at greater than 1% in at least two of ten replicate communities simulated in each condition. Simulations are based mostly on fitness differences estimated over a growth cycle in an environment containing a single limiting resource.

## Discussion

We quantified the fitness effect of six different ARGs in each of 11 host strains. Selection for genes depends on the net effect of positive selection for the function they encode and selection against any costs of expressing that function [50,51]. ARGs provide a benefit by protecting cells in the presence of a cognate antibiotic but many ARGs have been isolated from environments where antibiotics are at low concentrations and where costs might be influential in determining their fate [10,12–14].

Supporting the need to consider the role of ARG costs in determining their dynamics, we found that ARG costs were common (Figs 1, 2, S2). Considered over the two assay environments, all ARGs except *bla_SHV12_* were costly in at least one host. ARG effects differed over host strains, but the fitness of all tested hosts was reduced by at least one ARG (Fig 2, S2). Conversely, all ARGs were neutral or beneficial in at least one host strain, indicating the potential for specific strains to provide a ‘selective refuge’ for ARGs in the absence of antibiotic selection (Fig 2, S2).

Co-existence theory has been developed to understand the role of specific interactions in determining broader community dynamics [52–54]. This theory has recently been applied to consider ARG success given the availability of a range of environmental niches in which host strains have different fitness [11]. One key finding of that work was that ARGs that reduce host fitness in fewer environmental niches will generally be more successful than those that impose general costs over most environments. Because ARGs are frequently encoded by horizontally mobile elements that can transfer between hosts, different host strains can be thought of as distinct niches with transfer facilitating fluctuating selection. From the perspective of the ARG, strains in which they confer low costs represent refuges allowing them to minimize niche overlap with competitors. The more refuge strains that are available to an ARG, the more successful it is expected to be.

We have not directly tested the role of refuge strains on ARG persistence, but, even in the absence of any horizontal transfer, our simulations indicate that measured strain-dependent ARG fitness effects are of a magnitude that can influence the outcome of community composition (Fig. 3). We also note that ARG-strain interactions depended on the assay environment, a finding consistent with other studies measuring effects of antibiotic resistance [16,55–58] . This dependence increases the chance that refuge strains will be available for any given ARG but may also select for transfer between strains if costs change following environmental shifts.

We emphasise that the important result that strain-dependent ARG costs can influence strain and ARG success derive from a simple model that considers fitness estimates based on integrated growth over a competition cycle with a single limiting nutrient. We do not incorporate frequency dependence, and do not include direct inter-strain interactions, so that each starting community must eventually fix the single strain-ARG combination with the highest fitness. Consideration of more complex competitive interactions will increase the likelihood of diverse communities emerging, making it more likely that strain-dependent ARG effects will be relevant to determining final community compositions [59]. Unfortunately, competitive interactions among the strains we use are not well enough understood to be included in a model, but we are beginning community competition experiments to test directly test their influence.

Although the influence of strain-dependent ARG costs on ARG dynamics has not, to our knowledge, been examined directly, there are observations that are consistent with the idea that differential costs can affect ARG success. In one study trimethoprim use was decreased by 85% for two years in one Swedish county [60]. This change was correlated with a small overall decrease in frequency of the *dfrA* resistance gene but this small change masked larger changes within different *E. coli* subpopulations. Trimethoprim resistance was both unevenly distributed across subpopulations at the beginning of the intervention and changed differently over subpopulations during the intervention, decreasing from 67% to 41% in isolates from one subpopulation, but increasing from 17% to 31% in isolates from another [60].The authors note that explanations other than strain-specific differential fitness costs are possible, including clonal expansions of sequence types within subpopulations may influence ARG frequencies, though the frequent horizontal transfer of resistance genes means that they are unlikely to be the sole explanation.

One complication of predictions based on co-existence theory is that costs of ARGs can evolve. Indeed, costs of resistance phenotypes, especially those due to spontaneous mutations, are often reduced during growth in antibiotic-free environments (reviewed in 13,15). Several studies have also demonstrated changes in the fitness effect of ARGs due to chromosomal mutations [30,61,62]. A consequence of evolved changes in ARG costs is that ARG effects will not only differ across host stains and environments, but also through time as host strains adapt and compensate for any costs they confer. How this effect might influence long-term ARG fitness will be an important area for future study.

Finally, we note one example in our findings that highlights how difficult it is likely to be to predict ARG fitness costs. Within the set of six ARGs we considered, three—*bla*_TEM116*_, *bla*_CTX-M-15_ and *bla*_SHV12_—were β-lactamases. We expected the similar function of the encoded gene products to lead to similar host dependent effects, but in fact these genes showed a significant host strain interaction effect [63]. Indeed, one, *bla*_TEM116*_, conferred the highest cost of the genes we considered while another, *bla*_CTX-M-15_, conferred the highest benefit (Fig. 1). While we do not focus on the molecular basis of costs, we note that previous work on the effects of β-lactamases have proposed several mechanisms that may be relevant to our findings. β-lactamases need to be transported to the periplasmic region of cells to function and some signal peptides that direct this secretion have been associated with a fitness cost [20]. β-lactamases also have similarity with the penicillin binding proteins (PBPs) that process peptidoglycan components of cell walls. Residual activity of β- lactamases for PBP substrates might interfere with the dynamic process of peptidoglycan processing required to accommodate cell growth [21,22]. The host strains we used are genetically distinct, likely leading to different metabolic, physiological, and regulatory backgrounds on which effects of ARGs are tested. Whatever the mechanism of ARG costs, differences among our host strains evidently provide opportunity for them to play out differently.

Investigation of the influence of ARGs on microbial communities has been dominated by consideration of their effects in the presence of antibiotics. It was generally thought that ARGs would impose costs that would select against them in environments without this selection pressure, but it now seems that this assumption was optimistic [64]. While many ARGs can impose costs, a dependence on environment and host strain may create refuges that keep ARG frequencies high even in the absence of direct selection.

## Acknowledgements

This work was supported by a grant from the Royal Society of New Zealand Marsden Foundation (19-MAU-082 to T.F.C.). T.F.C. will make the strains constructed in this study available to qualified recipients following completion of an institutional material transfer agreement. The results of the competition experiments, summary input data, and analysis scripts that pertain to the experiments and analyses reported in this paper have been deposited at http://dx.doi.org/XX.XXX/dryad.XXXX.

## Notes

### Competing Interest Statement

The authors have declared no competing interest.

### Summary of Updates

Figures presenting resistance gene effects have been revised to increase clarity.

