## Supplementary Information for "Costs of antibiotic resistance genes depend on host strain and environment and can influence community composition"

Table S1. Source of ARGs used in this study.

| ARG | Primer pairs | Source of PCR template | Amplified region relative to CDS | |
|---|---|---|---|---|
| | | | upstream | downstream |
| *aadA* | ccgacgtctaagaaaccatgaattcctactcgagttcatgtgcagc<br>cctcgtgatacgcctatttctgacgctcagtggaacg | pTargetF plasmid (Addgene, #62226) | 100* | 33* |
| *catI* | ccgacgtctaagaaaccatgaattcttctggtcggcgcatagctg<br>cctcgtgatacgcctattgcaagcagcagattacgcg | portMAGE4 plasmid (Addgene, #72679) | 178 | 25 |
| *dfrA5* | ccgacgtctaagaaaccatgaattccatggcttgttatgactg<br>cctcgtgatacgcctatttgtgcttagtgcatctaacg | *E. coli* strain B706 | 158 | 93 |
| *bla<sub>TEM-116</sub>*†* | ccgacgtctaagaaaccatgaattcttctggtcggcgcatagctg<br>cctcgtgatacgcctattgcaagcagcagattacgcg | portMAGE2 plasmid (Addgene, #72677) | 72 | 147 |
| *bla<sub>CTX-M-15</sub>* | ccgacgtctaagaaaccatgaattc<br>cctcgtgatacgcctatt | Sequence was synthesised based on GenBank: MK125035.1. | 250 | 150 |
| *bla<sub>SHV-12</sub>* | ccgacgtctaagaaaccatgaattc<br>cctcgtgatacgcctatt | Sequence was synthesised based on GenBank: CP048293.1. | 237 | 150 |

* The upstream and downstream regions of *aadA* on pTargetF were contributed by aminoglycoside N-acetyltransferase AAC(3)-Iva (AAC(3)-IV5a).
†The portMAGE4 encoded BlaTEM (BlaTEM-116*) has more similarity to BlaTEM-116 than BlaTEM-1, but has one amino acid difference from BlaTEM-116 at amino acid 274 (Q in BlaTEM-116 but R in BlaTEM-116*). BlaTEM-116 differs from BlaTEM-1 at amino acid 82 and 184 (I and A respectively in BlaTEM-1, but V and V in BlaTEM-116).

Table S2. Phylogenetic signal of ARG fitness effects in antibiotic-free environment.

| | ARG | Pagel's lambda* | Blomberg's K* |
|---|---|---|---|
| Core gene phylogeny | *aadA* | 0.44 | 0.06 |
| | *bla*$_{CTX-M-15}$ | <0.001 | 0.02 |
| | *bla*$_{SHV12}$ | 0.15 | 0.03 |
| | *bla*$_{TEM116*}$ | 0.99 | 0.48 |
| | *cat* | 0.30 | 0.14 |
| | *dfrA5* | <0.001 | 0.15 |
| Accessory gene phylogeny | *aadA* | 0.62 | 0.41 |
| | *bla*$_{CTX-M-15}$ | <0.001 | 0.14 |
| | *bla*$_{SHV12}$ | 0.26 | 0.25 |
| | *bla*$_{TEM116*}$ | <0.001 | 0.61 |
| | *cat* | 0.31 | 0.50 |
| | *dfrA5* | <0.001 | 0.44 |

*Pagel's lambda and Blomberg's K metrics assume a Brownian motion model of trait evolution. If traits are phylogenetically independent both metrics give a value close to 0. In no case was the estimated metric significant at $P < 0.05$, indicating a lack of phylogenetic signal.

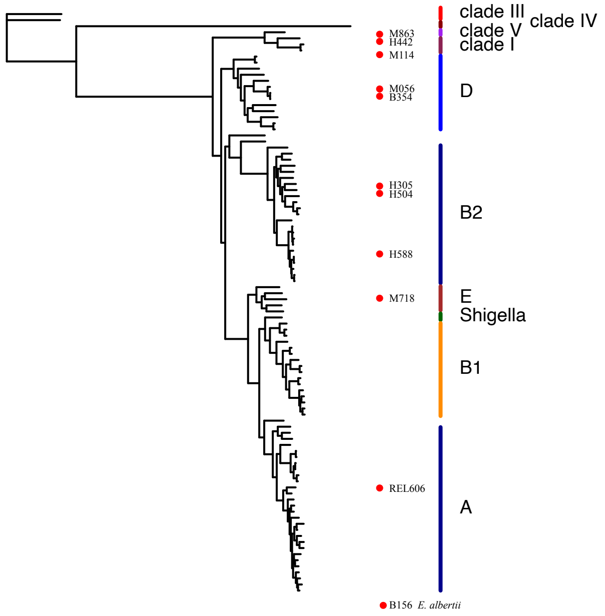


Figure S1. Maximum likelihood core-genome phylogeny of a set of 96 diverse *Escherichia* spp. indicating strains used in this study. Coloured lines and associated labels indicate recognized clades. Strain B156 is classified as *E. albertii* and is not included in our phylogenetic analysis. Details in Wang et al. Proc. Natl. Acad. Sci. USA 113: 5047-5052.

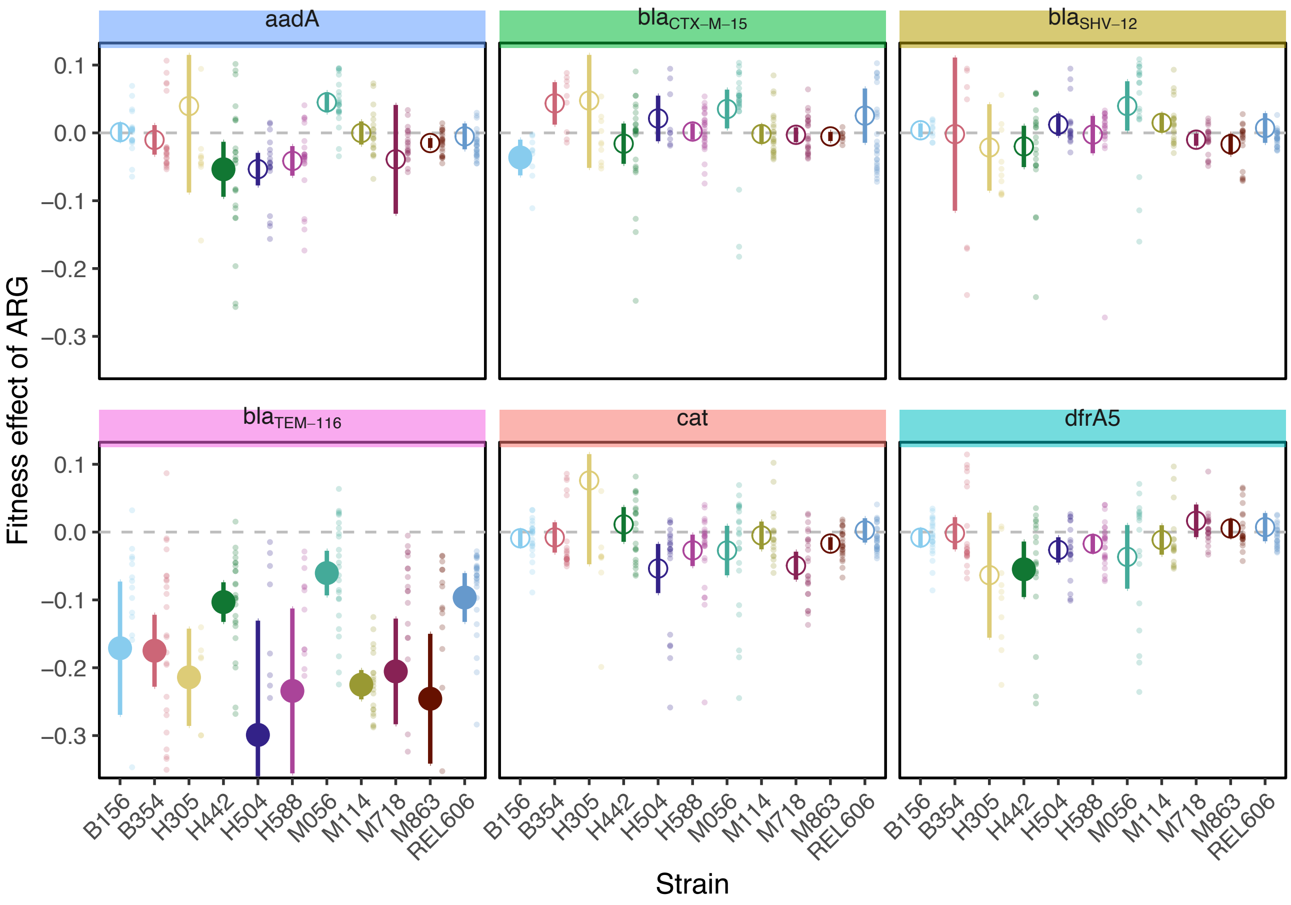


**Figure S2. Fitness effect of ARGs in each host strain measured in an environment containing an unrelated antibiotic.** For each host-ARG combination large symbols indicate mean fitness over all replicate estimates and error bars indicate 95% CI. Fitness assays were carried out in an environment containing kanamycin, to which all strains are resistant through carriage of a resistance gene encoded separately to the six focal genes. Large symbols are filled if the mean ARG effect is different from 0 after correction from multiple comparisons using Dunnett's test (M&M for details). Background points indicate individual estimates used to calculate each mean. The horizontal dashed line indicates no fitness effect. Background label panels are coloured to match the ARG color used in figure 1. n ≥ 5 for each host-ARG combination.
